# Impact of in vivo cyclic reprogramming on the choroid plexus

**DOI:** 10.1101/2023.02.28.530533

**Authors:** Jessica Avila Lopez, Clauda Abboud, Maged Ibrahim, Javier Rocha Ahumada, Mariano Avino, Mélanie Plourde, Karl Fernandes, C. Florian Bentzinger, Benoit Laurent

**Affiliations:** Research Center on Aging, Centre Intégré Universitaire de Santé et Services Sociaux de l’Estrie-Centre Hospitalier Universitaire de Sherbrooke, Sherbrooke, QC, Canada; Department of Biochemistry and Functional Genomics, Faculty of Medicine and Health Sciences, Université de Sherbrooke, Sherbrooke, QC, Canada; Département de Pharmacologie-Physiologie, Faculté de Médecine et des Sciences de la Santé, Institut de Pharmacologie de Sherbrooke, Centre de Recherche du Centre Hospitalier Universitaire de Sherbrooke, Université de Sherbrooke, Sherbrooke, QC, Canada; Bio-informatic platform, Department of Biochemistry and Functional Genomics, Faculty of Medicine and Health Sciences, Université de Sherbrooke, Sherbrooke, QC, Canada; Department of Medicine, Faculty of Medicine and Health Sciences, Université de Sherbrooke, Sherbrooke, QC, Canada

## Abstract

In vivo reprogramming using the transient expression of Oct3/4, Sox2, Klf4 and c□Myc (OSKM) transcription factors can be used to induce tissue regeneration. A cyclic regime for short□term OSKM expression has been shown to promote regeneration of several organs however its impact on the brain remains largely unknown. We investigated the effects of a cyclic short-term OSKM expression on the choroid plexus (CP), a highly vascularized tissue found within the brain ventricles which is responsible for producing the cerebrospinal fluid (CSF). Transient reprogramming was done on 8-week-old mice carrying the polycistronic OSKM cassette under tetracycline operator (tetO) and confirmed the successful transient reprogramming. We then performed the analysis of the CP at cellular and molecular levels. The CP tissue exhibited minor morphological changes in height and area of epithelial cells. We did not observe any significant differences in the integrity of the brain-CSF barrier but noticed an increase of NKCC1 expression, a protein involved in CSF production. A whole transcriptome analysis (RNA-seq) was also carried on the tissue and showed no difference in gene expression after the transient reprogramming, at the exception of blood-related genes. Our results indicate that surprisingly the CP mainly remains insensible to in vivo transient reprogramming as only morphological and protein changes were observed in the tissue, suggesting that translational changes might be at stake during the reprogramming process but are not present at the transcriptomic level. Our results also highlight that more tailored strategies need to be developed for exploring the potential of CP reprogramming in regenerative medicine.

## Introduction

In the last decades, life expectancy has significantly improved going from 46 years old in 1950 to 70 years old in 2019 (Roser et al., 2013). However, the extension of lifespan has also increased the risks of developing age-related diseases. Indeed, aging represents an accumulation of time-dependent biological and physiological changes over time (López-Otín et al., 2013). Indeed, many studies have described morphological, physiological and metabolic modifications occurring in the aging brain, as reviewed by Mattson and Arymygam (2018) and it is also the primary risk factor for the development of neurodegenerative diseases (ND) (Hou et al., 2019).

For years the human specie have seeked different methods that enable the rejuvenation of different cells and organisms with the hope to extrapolate it to the humankind someday (de Magalhães & Ocampo, 2022). Gurdon (1962) was the pioneer in reprogramming somatic cells to pluriopotent ones in an embrionic state using the somatic cell nuclear transfer (SCNT) technique. The SCNT allowed (Wilmut et al., 1997) to generate the first mammalian generated by somatic cloning: sheep Dolly. These two discoveries proved that the somatic cells contained all the needed genetic information to generate a new organism and also stated that the egg cell environment contained the needed factors to do so. In 2001 (Tada et al., 2001) manage to fuse somatic cells with embryonic stem cells (ESCs) generating cells with puripotency gene expression but they had a limited supply and could generate immune rejection when implated to living organisms plus the technic had many ethical concerns. The induced Pluripotent Stem Cells (iPSCs) developed by (Takahashi & Yamanaka, 2006) arrived as an alternative to solve the problems the ESCs had. They seeked for the factors that give the ESCs their pluripotency, identifying 24 of them out from which 4 where described as essential to promote the reprogramming of somatic cells to “embrionic stem cell-like” cells: Oct4, Sox2, Klf4 and c-Myc (OSKM) (Takahashi & Yamanaka, 2006).

By promoting the expression of OSKM different teams have proved there is reprogramming in different organs in-vivo such as in the liver (Chondronasiou et al., 2022; Hishida et al., 2022), pancreas (Chondronasiou et al., 2022), spleen (Chondronasiou et al., 2022; Ocampo et al., 2016), skin (Doeser et al., 2018; Ocampo et al., 2016) and Skeletal muscle (Ocampo et al., 2016). When it comes to the CNS reprogramming have been observed in vitro in the neural stem cells (Han et al., 2021; Yao et al., 2015) and in vivo specifically in the Dentate Gyrus Cells (DG) (Rodríguez-Matellán et al., 2020) but the reprogramming of the Choroid Plexus (CP) has not been described.

The CP is a monolayer of epithelial cells located in each brain ventricle giving a total of four CPs that interacts with both the Cerebrospinal Fluid (CSF) and blood forming a physical barrier called the Blood-CSF-Barrier (BCSFB). There are three major fluids linked to the central Nervous system (CNS): the blood, the Interstitial Fluid (ISF) and the cerebrospinal fluid (CSF). The ISF surrounds the parenchymal cells and the spinal cord (Weller et al., 2010). It promotes an environment for cells to survive and function normally. Its ion concentration is important for controlling the nervous activity and its osmolarity regulate cell volume (Hladky & Barrand, 2014). The CSF is the liquid within the brain ventricles responsible to drain the metabolites out of the brain, performs the osmoregulation of inorganic ions, buffers the pH and transports micronutrients (Ghersi-Egea et al., 2018) therefore a key element to look at for markers of brain function dysregulation (Scheiblich et al., 2020). The CSF is produced by the CP (Dato et al., 2017; Javed et al., 2022) on a highly vascularized stroma with fenestrated capillaries.

During aging, morphological changes of the CP have been observed such as: the Choroid Plexus Epithelial cells (CPECs) decrease in height by 11% in humans, 15% in rat animal models and 9.5% in mice animal model; the nucleus losses its round shape and it flattens; the microvilli on CPECs are shorter (Serot et al., 2000; Avila Lopez, 2022); the stroma thickness increases with age, with the appearance of fibrosis (Serot et al., 2001); the epithelial basal membranes thickened up (Avila Lopez, 2022; Serot et al., 2001). Appart from producing the CSF and forming the BCSFB, the CP plays an important role in the maintenance of the brain homeostasis by cleansing brain metabolites and secreting important factors (Van Cauwenberghe et al., 2020). The CPECs are not renewed through the lifespan (McDonald & Green, 1987) compromising the CP well-functioning during the lifespan. The ability to reprogram the brain, specifically the CP could prevent the brain’s morphological, physiological and metobolic deterioration.

## Material and Methods

### Mice

All experiments were approved by the animal care and use committee of the Université de Sherbrooke under the protocol number 2018-2132. Mice with the polycistronic OSKM cassette (Oct4, Sox2, Klf4, c-Myc) and the rtTA transactivator (R26-rtTA;tetO-OSKM) were obtained from The Jackson Laboratory (Cat. No023749). All experiments were performed using 8 week-old heterozygous male and female mice. Induction of OSKM was performed by administration of doxycycline (Dox, 1mg/ml, Cat. No D9891, Sigma-Aldrich) in the drinking water of mice supplemented with 5% sucrose (Cat. No 97061-428, VWR international). All mice were treated for 3 days with Dox followed by 4 days of withdrawal. All mice were bred, housed in standard cages under specific pathogen-free conditions and were fed standard mouse chow and water ad libitum and kept on a 12h light/dark cycle.

### Skeletal muscle injury

Mice were anesthetized with isoflurane (Cat. No 0-0-12593, Pharmacie de l’Hôpital Fleurimont) and the tibialis anterior (TA) muscle was injured by intramuscular injection of 50μL of cardiotoxin (12μM) from Naja Pallida (Cat. No L8102, Latoxan). Following injury, mice were treated once with buprenorphine (0.05-1mg/Kg, McGill University).

### Quantitative PCR

Total RNA was extracted from muscles using Ribozol (Cat. No CA97064-950, VWR international) according to the manufacturer’s protocol. The A260/280 ratio of the isolated RNA was 1.8-2.0. mRNA was reverse transcribed into single-stranded cDNA using the SensiFAST cDNA Synthesis Kit for RT-PCR (Cat. No BIO-65054, FroggaBio). After conversion to cDNA, real-time PCR was performed using the SensiFAST SYBR No-ROX Kit (Cat. No BIO-98005, FroggaBio) using a Rotor-Gene 6000 (Corbett Life Science). PCR primers for Sox2 and the housekeeper mATP5B were derived from (Lukjanenko et al., 2016, 2019; Ocampo et al., 2016).

### RNA-Seq library preparation

The RNA isolation from the LVCP was performed with the NucleoSpin RNA plus XQ (Takara) following the manusfacturer’s protocol. RNA-seq libraries were prepared using NEBNext® single cell/low input RNA library preparation Kit (New England Biolabs). Library preparation was initiated with 10ng of total RNA. RNA along with all the other reagents were thawed on ice. Further steps were conducted according to the manufacturer’s protocol until the final library amplification. Each final library was then purified using the MinElute PCR Purification Kit (Qiagen) and then quantified on a Qubit system (Thermo Fisher Scientific). Multiplexed libraries were pooled in approximately equimolar ratios and were then run in a 2% agarose gel. DNA between 300 and 700 base pairs was then cut and purified using the Qiagen Gel Extraction Kit (Qiagen). Library sequencing was performed at the RNomics platform of the Université de Sherbrooke (Canada). Single-End sequencing was performed on all the samples with a 75bp sequencing depth on a NextSeq machine.

### RNA-Seq Analysis

FastQC toolkit (version 0.11.8) with the default parameters to filter the low-quality reads was used.

TrimGalore (version 0.6.4) was set to pair reads with the default parameters. Was used to trim the reads, N was set to, 5 and end clips were set to 3. STAR software (version 2.7.3a) with the default parameters, but for the maximal number of multi mapping locations which was set to 10 and SAM primary flag was set to all best scores, was used to map the reads to the mouse genome (GRCm38, GENCODE vM23, primary assembly). The evaluation of gene expression (CPM) was done by FeatureCounts from the SubRead package (version 2.0.0). Exons were set as features and grouped by gene. R (version 3.5.3) was used for the statistical analysis using the edgeR package (version 3.24.3) and using python as well (version 3.7.3). Significant change in gene expression (FDR of 1%) was done by a decision tree-based-recursive-binary-split-algorithm which was built using scikit-learn package (version 0.23.1) in python.

### Histological analysis

The evaluated organs were washed in PBS 1x and placed into an histology cassette. The cassette was submerged in PFA 4% diluted in PBS 1x, in a volume of PFA 4% 10 times greater than the volume of the tissue. The cassette rested in the PFA 4% for 16 hours at 4 °C. The fixed brain tissue was then washed three times in ethanol 70%. The cassette was embedded in paraffin. The bloc was cut every 4 microns and two consecutive cuts were placed on a glass slide for its further analysis.

### Morphology analysis

The slides were stained with Hematoxylin and Eosin (H&E) for the morphology analysis following a modified protocol from Cardiff et al., (2014). The Hematoxylin (Fisher, catalog number: 23245651) was used pure, while was used a 1:3 solution of Eosin (Fisher, catalog number: 23245827) and EtOH 95%/ 0.3% acetic acid. The acid water was prepared by diluting 12 drops of HCL 20% in 750 ml of distilled water, the basic water was prepared by adding 16 drops of ammonium hydroxide in 750 ml of distilled water. The morphology analysis was performed by scanning the slides in the Nanozoomer of Hamamatsu and then in the software NDP.view2 (Hamamatsu) software with the brain slides.

## Results

### In vivo cyclic reprogramming

Our cyclic reprogramming paradigm consisted in the induction of the OSKM factors by the administration of doxycycline in the drinking water supplemented with 5% sucrose where heterozygous male and female mice of 8 week-old were treated for 3 days with Dox followed by 4 days of withdrawal, at day 16 an injury in the tibialis anterior (TA) muscle by the intramuscular injection of 50μL of cardiotoxin (12μM) was done in some mice from the R26-rtTa;+ and R26-rtTa;tetO-OSKM groups (***Figure 1A***). Small regenerating fibers were observed in the skeletal muscle of R26-rtTa;tetO-OSKM (***Figure 1B***) indicating a reprogramming process at stake. To verify the presence of the Yamanaka factor in the skeletal muscle we performed a qPCR of this tissue for SOX2 which was significantly increased in the R26-rtTa;tetO-OSKM group when compared to R26-rtTa;+ (***Figure 1C***).

**Figure 1.**
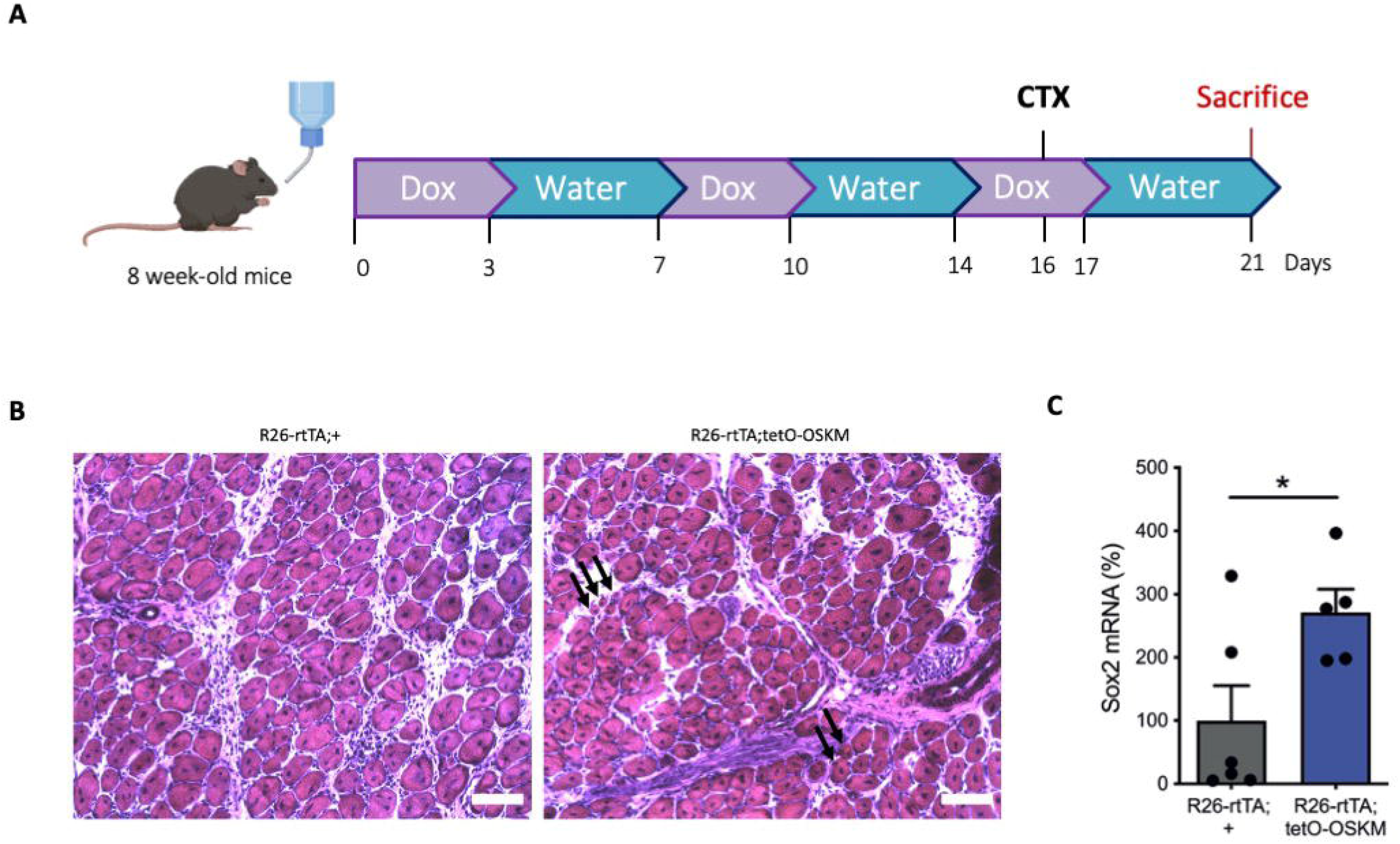
Cyclic reprogramming paradigm. **(A)** Graphical outline of the 3-week cyclic treatment protocol with the timepoint of cardiotoxin (CTX) injury. In details, 8-week-old mice achieved three cycles of treatment consisting of drinking water with doxycycline (Dox) for 3 days followed water without Dox for 4 days. At the end of these 3 cycles, the sacrifice of mice was performed. **(B)** Representative hematoxylin and eosin staining of skeletal muscle cross sections of R26-rtTA;tetO-OSKM mice and R26-rtTA;+ control mice at 5 days post-CTX injury after three cyclic Dox treatments. Black arrows indicate accumulation of groups of small regenerating fibers. Scale bar: 40 *μ*m. **(C)** Expression of the polycistronic OSKM cassette in skeletal muscle assessed by quantitative PCR (qPCR). mRNA levels of Sox2 were measured in R26-rtTA;tetO-OSKM mice (n=5) and R26-rtTA;+ control mice (n=6) at day 21, and normalized to that of ATP5B. Results are indicated as mean ± SD. (*) p < 0.05 from unpaired t-test with equal means.

One hallmark of a successful systemic reprogramming is the appearance of teratomas in different peripheral organs such as the liver, kidney and the spleen. Therefore, we decided to collect these organs and evaluate the morphological changes between both groups. We observed that all R26-rtTa;+ individuals did not have teratomas while all R26-rtTa;tetO-OSKM samples (n=4) but one had teratomas (***Figure 2A***) This information confirms the presence of the OSKM factors in the liver, kidney and spleen too. A significant increase in the number of teratomas per liver section evaluated was observed in the R26-rtTa;tetO-OSKM group (***Figure 2B***). A greater area is occupied by teratomas in the liver of R26-rtTA;tetO-OSKM mice after three cyclic Dox treatments (***Figure 2C***).

**Figure 2.**
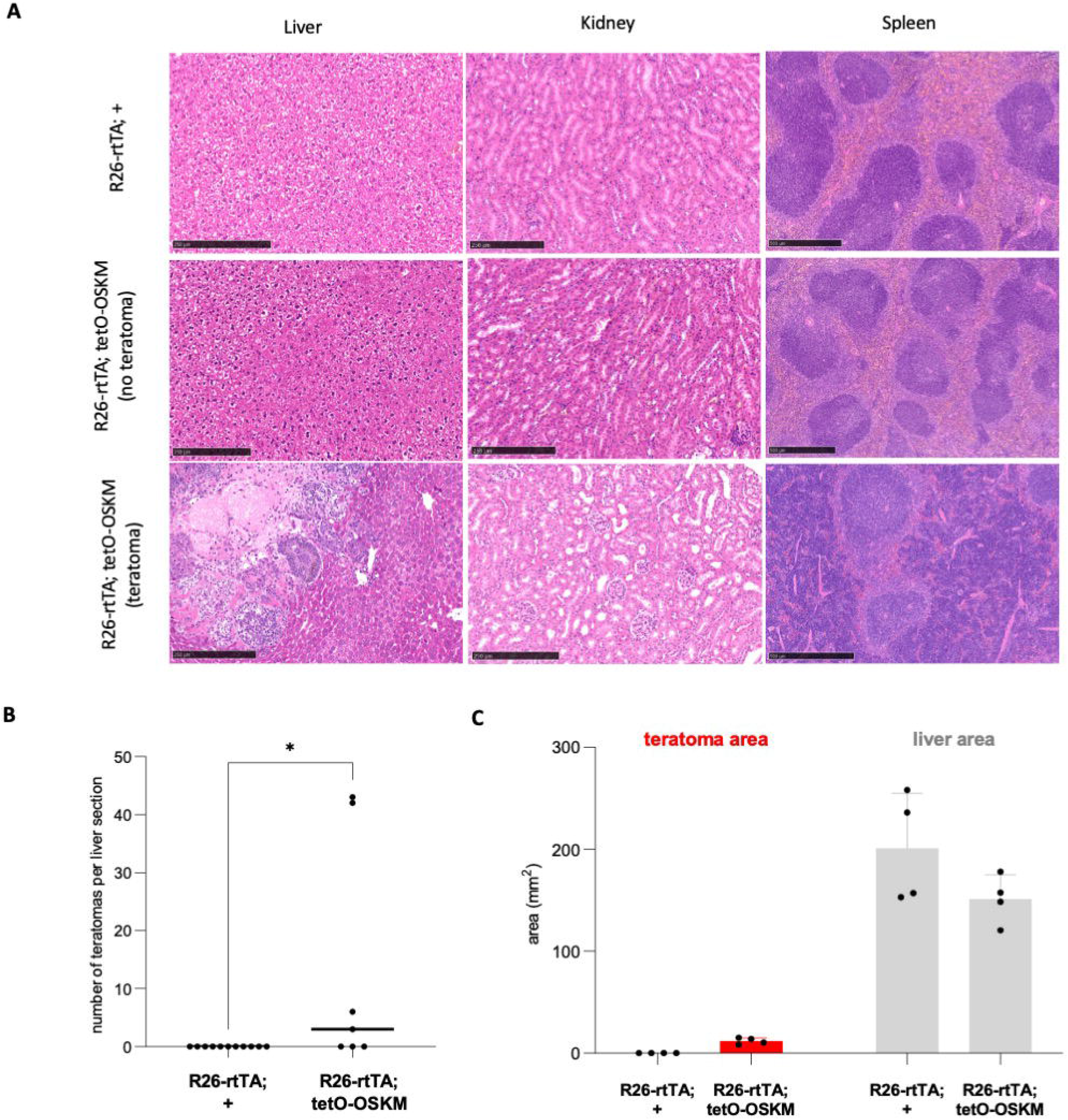
Effects of global cyclic OSKM induction. **(A)** Representative hematoxylin and eosin staining of liver, kidney and spleen cross sections of R26-rtTA;+ control mice (top panel) and R26-rtTA;tetO-OSKM mice (middle and bottom panels) after three cyclic Dox treatments. Scale bar: 250 *μ*m. **(B)** Number of teratomas per liver section of R26-rtTA;tetO-OSKM mice (n=7) and R26-rtTA;+ control mice (n=11) after three cyclic Dox treatments (day 21). **(C)** Area occupied by teratomas in the liver of R26-rtTA;tetO-OSKM mice (n=4) and R26-rtTA;+ control mice (n=4) after three cyclic Dox treatments.

### Morphological changes in the brain and choroid plexus

Teratomas were found in the brain of 3 out of 4 R26-rtTA;tetO-OSKM mice whereas none of the R26-rtTA;+ mice had any teratomas (***Figure 3A***). Morphological changes of the lateral ventricle choroid plexus (LVCP) were found (***Figure 3B***). There is a significant increase in the height of the LVCP epithelial cells of the R26-rtTA;tetO-OSKM mice (***Figure 3C***) and so in their area (***Figure 3D***).

**Figure 3.**
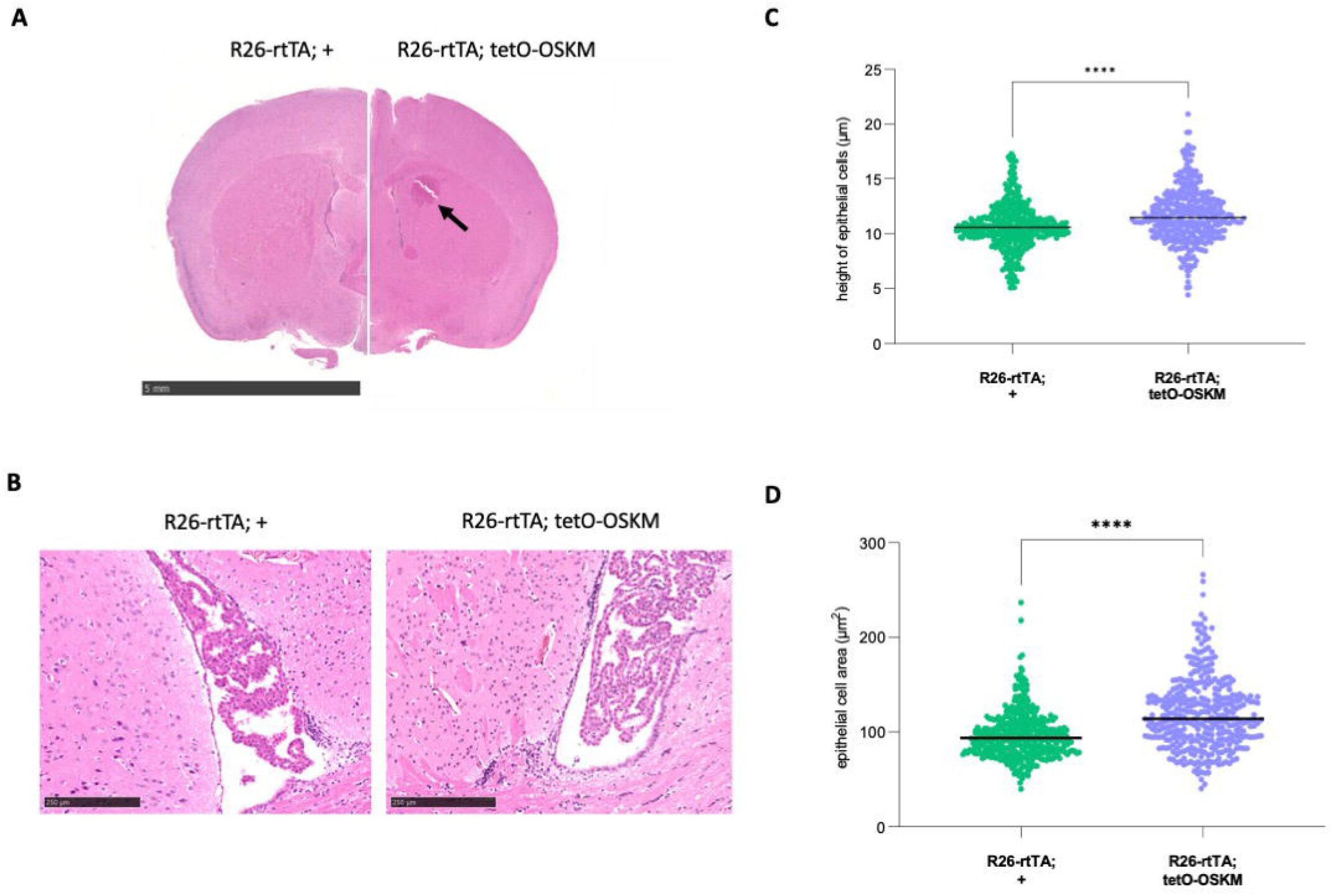
Effects of global cyclic OSKM induction on the brain and choroid plexus. **(A)** Representative coronal cuts of whole brain of R26-rtTA;tetO-OSKM mice (right panel) and R26-rtTA;+ control mice (left panel) stained with hematoxylin and eosin. Teratoma is indicated with a black arrow. Scale bar: 5 mm. **(B)** Representative coronal cuts of the ventricular space in which the plexus choroid is located. Cuts were stained with hematoxylin and eosin. Scale bar: 250 *μ*m **(C-D)** Morphological changes of choroid plexus epithelial cells were measured after three cyclic Dox treatments (day 21). The height of epithelial cells **(C)** and epithelial cell area **(D)** were measured for the choroid plexus of of R26-rtTA;tetO-OSKM mice and R26-rtTA;+ control mice (n=400 counts per group). T-test with Welch’s correction. *** P≤ 0.0001

### No major transcriptomic changes occur in the CP

Knowing that the LVCP morphology changes between groups, we decided to evaluate its transcriptomic changes by performing a bulk RNA-seq. No major transcriptomic changes were observed when comparing the expression of 27,164 genes in the LVCP of R26-rtTA;+ control mice and R26-rtTA;tetO-OSKM mice (n=3 per group) (***Figure 4A***). Only 16 genes were differentially expressed (FDR<0.05) (***Figure 4B***) among which four blood-related genes were significantly differentially expressed in the R26-rtTA;tetO-OSKM group (*Hba-a1, Hbb-b1, Hba-a2 and Hbb-bs*) (***Figure 4C***). A list of the 25 most abundantly expressed genes annotated to be secreted factors by the LVCP described by (Lun et al., 2015) was done but only the *Bgn* gene was differentially expressed between groups (***Figure 4D***).

**Figure 4.**
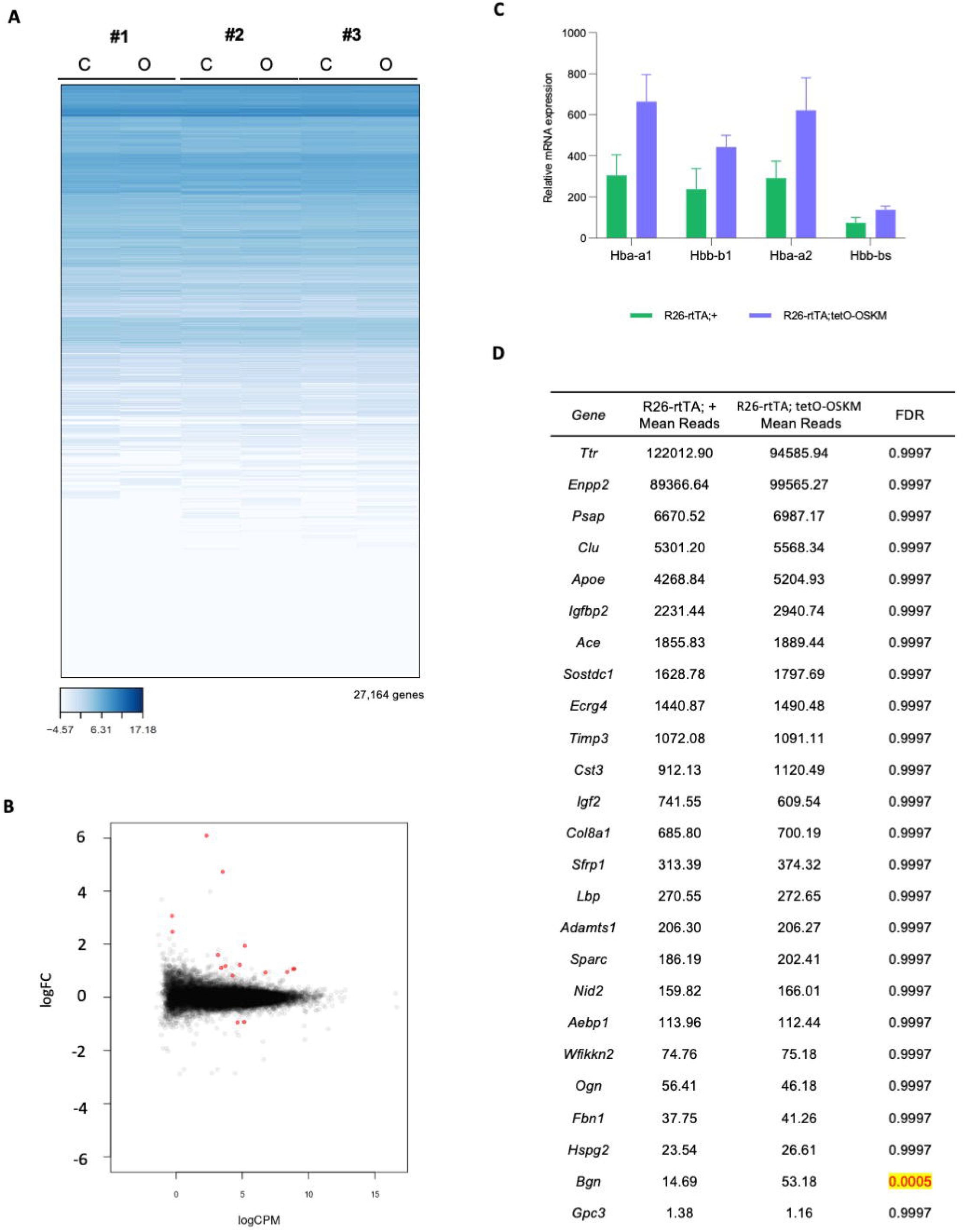
Transcriptome signature of the choroid plexus in response to OSKM induction. **(A)** Heatmap of the expression of 27,164 genes in the choroid plexuses of R26-rtTA;+ control mice (C) and R26-rtTA;tetO-OSKM mice (O) (n=3 per group). (#) indicates sibling pairs. **(B)** Smear plots of differentially expressed genes in the choroid plexus of R26-rtTA;tetO-OSKM mice. Differentially expressed genes compared with control were labelled as red points whereas insignificant genes were labelled as black. The mean expression level (logCPM) and the Fold Change of differentially expressed genes (logFC) are displayed. **(C)** Relative gene expression levels of blood-related differentially expressed genes in the choroid plexus of R26-rtTA;+ control mice and R26-rtTA;tetO-OSKM mice (n=3 per group). **(D)** Table of the 25 most abundantly expressed genes annotated to be secreted factors with the mean normalized read counts and FDR values.

### Translational changes occur in the CP after the reprogramming induction

The production of the CSF decreases by up to a 45% with aging and the changes in the morphology of the CP have been linked to this outcome (Damkier et al., 2013; May et al., 1990; Serot et al., 2001, 2003). As the only changes observed in the LVCP were in the morphological level but not the transcriptomic one, we decided to evaluate the expression of the NKCC1 protein (***Figure 5A***). which is a Na-K-Cl cotransporter responsible for 50% of CSF production (Steffensen et al., 2018). A significant increase in NKCC1 protein expression was observed in the R26-rtTA;tetO-OSKM subjects (***Figure 5B***). which is consistent to the increase of the cell height and area observed in the same individuals (***Figure3C-D***). The relative mRNA enrichment of the NKCC1 gene obtained from the bulk RNA-seq shows no difference between the two studied groups (***Figure 5C***).

**Figure 5.**
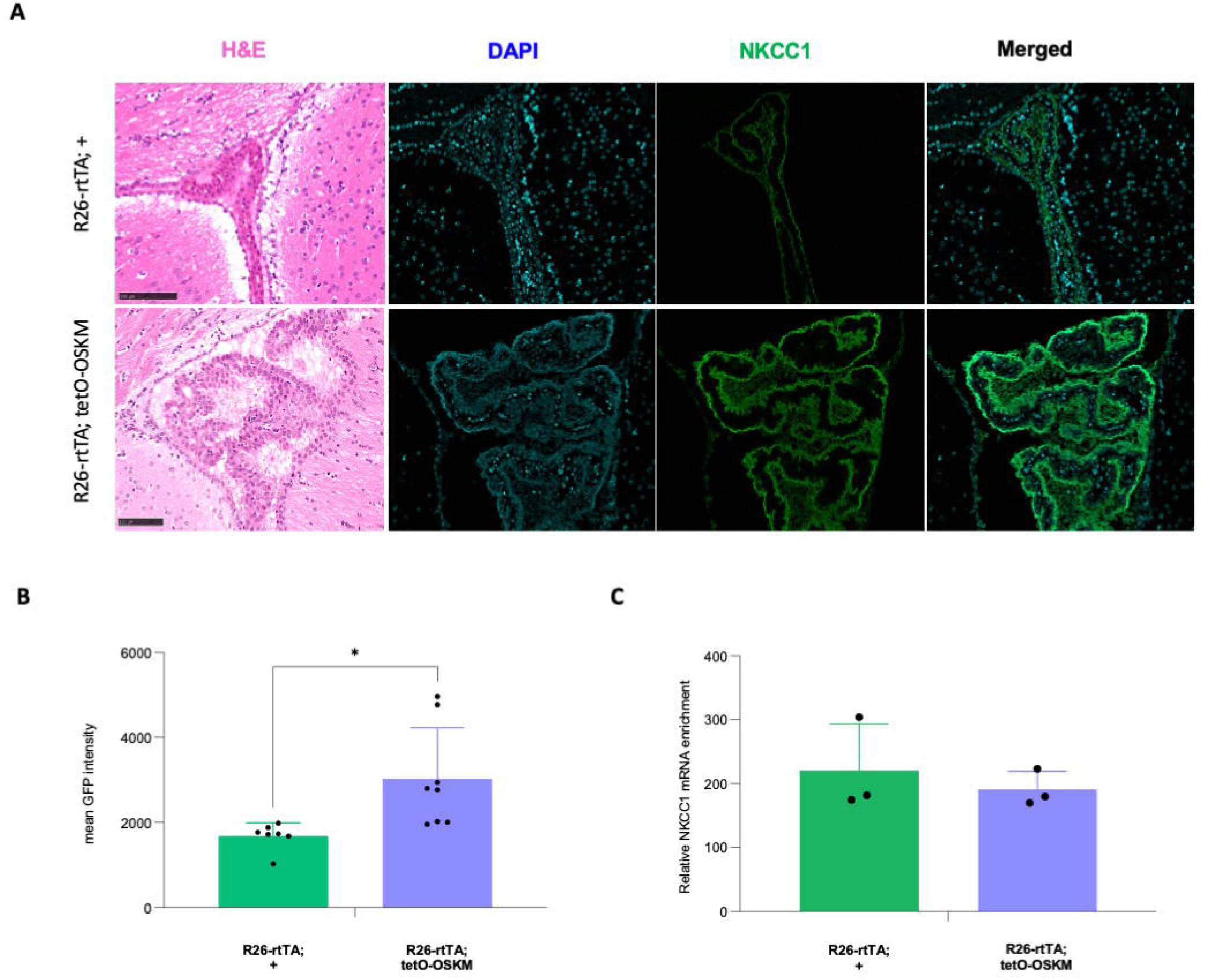
Analysis of NKCC1 expression in the choroid plexus. **(A)** NKCC1 immunofluorescence on coronal brain cuts of R26-rtTA;+ control mice and R26-rtTA;tetO-OSKM mice. Images illustrate the representative staining in the choroid plexus area with the NKCC1 antibody (green) and DAPI (blue). The left panel represents the hematoxylin and eosin staining, and the right image panel constitutes a merged image of NKCC1 and DAPI signals. Scale bar: 100 *μ*m. **(B)** Quantitative comparative analysis of NKCC1 expression in choroid plexus of R26-rtTA;+ control mice (n=7) and R26-rtTA;tetO-OSKM mice (n=8). (*) p < 0.05 from unpaired t-test for equal means. **(C)** Relative mRNA levels of NKCC1 in choroid plexus of R26-rtTA;+ control mice and R26-rtTA;tetO-OSKM mice (n=3 per group). These results were obtained in our transcriptomic analysis.

### Changes in the subventricular stem cell niche after the reprogramming induction

After observing an increase in the NKCC1 protein expression in the LVCP of the R26-rtTA;tetO-OSKM samples we decided to evaluate the expression of SOX2 in the subventricular stem cell niche as it is nourished directly by the CSF secreted from the LVCP which contains neurotrophic factors that promote the stem cell colony formation and proliferation within the brain (Silva-Vargas et al., 2016). We observed an increase in SOX2 expression in the reprogrammed mice (***Figure 6***).

**Figure 6.**
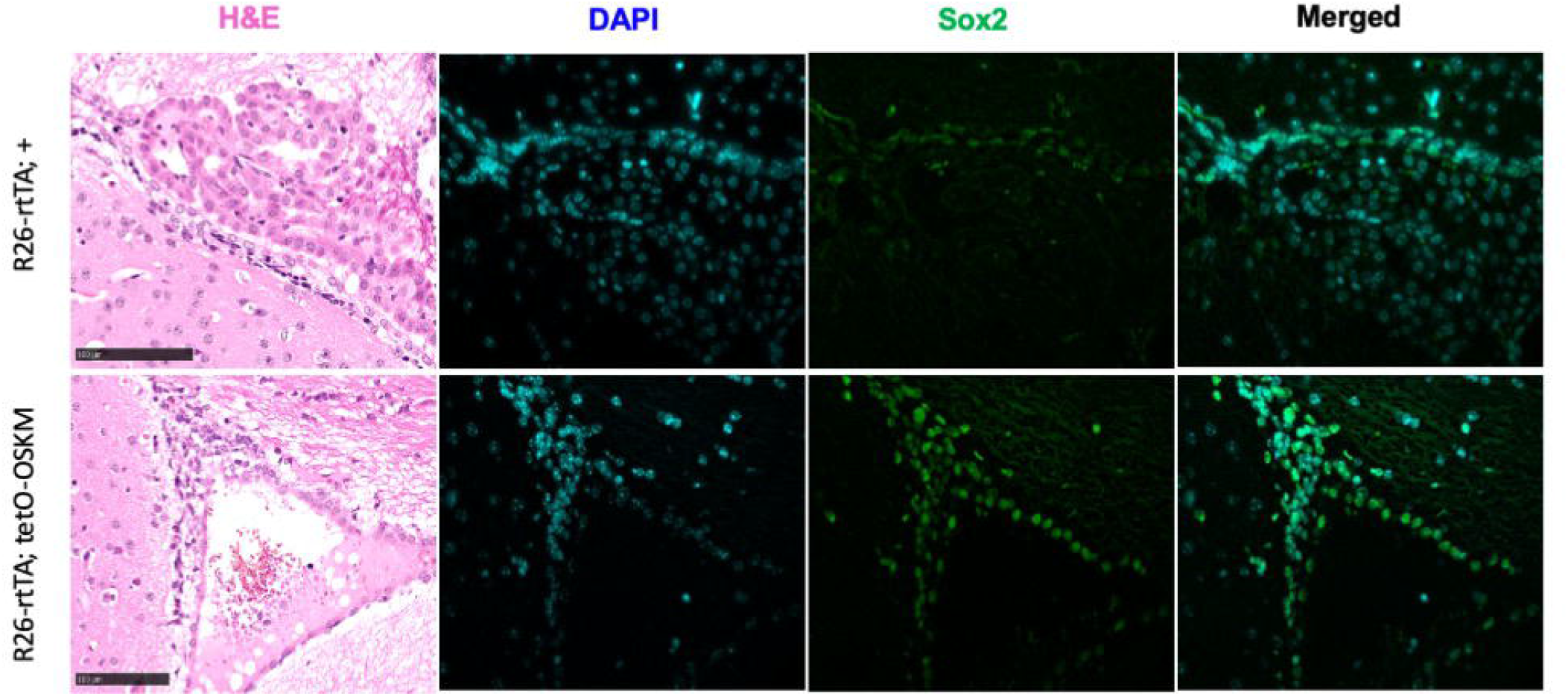
Analysis of Sox2 expression in the subventricular zone. Sox2 immunofluorescence on coronal brain cuts of R26-rtTA;+ control mice and R26-rtTA;tetO-OSKM mice. Images illustrate the representative staining in the choroid plexus area with the Sox2 antibody (green) and DAPI (blue). The left panel represents the hematoxylin and eosin staining, and the right image panel constitutes a merged image of Sox2 and DAPI signals. Scale bar: 100 *μ*m.

## Discussion

The evaluation of in-vivo induction of the OSKM factors in a mice model have been challenging as their survival rate depends on three different factors: the genotype of the mice, the dosage of Dox and the timeline of the induction. It have been proven that homozygous mice have a shorter life span (15 weeks) when compared to the heterozygous ones (35 weeks old) (Alle et al., 2022). The mice we worked with were heterozygous so we were expecting to have a higher survival rate than the described with the homozygous. The Dox dosage we gave was 1mg/ml as previously done by (Browder et al., 2022; Ocampo et al., 2016; Rodríguez-Matellán et al., 2020) but this dosage has been described by (Abad et al., 2013) in not convenient as it results in high morbidity after 1 week of induction suggesting to use smaller dosages of Dox such as 0.2mg/ml. The last factor to consider in the type of induction between a continuous or cyclic cycle. The continuous cycle promotes excessive teratomas in different peripheral organs such as liver and kidneys and leads to premature death (Abad et al., 2013; Ocampo et al., 2016). Therefore we used a cyclic OSKM induction protocol to reduce the high levels of mortality (Ocampo et al., 2016). Even when the Dox dosage could be considered as high, by working with heterozygous mice under a cyclic cycle of induction we considered our model as viable to further evaluate the effects of reprogramming in some peripheral organs such as skeletal muscle, liver, kidneys and spleen. Accumulation of groups of small regenerating fibers in the TA 5 days post-CTX injury after three cyclic Dox treatments was observed indicating the expression of the OSKM factors in the skeletal muscle as previously described by (Wang et al., 2021).

(Ocampo et al., 2016) regarding the presence of teratomas in only the homozygous OSKM mice. We found teratomas present in the liver, kidney, spleen and brain in our heterozygous animals too. Ocampo used the same concentration of Dox (1mg/ml) for the OSKM induction, the only difference is that the OSKM factor in his mice had the polycistronic cassette placed in the Col1a1 locus whereas ours had it in the Neto2 gene. Neto2 gene overexpression has been correlated with tumor progression in different studies (Fedorova et al., 2017; Hu et al., 2015). A better standardization in the locus where the OSKM factors are induced need to be established beforehand so results could be comparable. Parras (et al., 2022) have identified that the hepatical and intestinal failure is the reason of premature death in in vivo reprogramming models. They found this out by evaluating and comparing models where the OSKM factors where at the *Col1a1* and *Pparg* genes, they created a new reprogramming mouse strain where the OSKM factors are not expressed in the liver and small intestine where they saw a reduction in teratomas and a longer life in the mouse. For future experiments involving reprogramming in-vivo models the design of strains that don’t express the OSKM factor in liver and small intestine could lead to better efficiency.

During aging, morphological changes of the epithelial cells of the LVCP have been annotated such as a significant decrease in their height and area (Serot et al., 2000, 2001, 2003). We observed an increase in both the height and the cell area of the LVCP which increases the surface area available for the interaction between the stroma, which is the connective tissue that gives support to the cells (De Wever & Mareel, 2003) and to the CSF (Ghersi-Egea et al., 2018). Further analysis of the mechanisms at stake to increase the height and area of the LVCP cells need to be done as a possible therapeutic avenue for neurodegenerative disease that show to have a decrease such as in Alzheimer’s disease (Kaur et al., 2016).

Previous research teams have proven the ability of the doxycycline to trespass through the blood-CSF-barrier (BCSFB) (Nau et al., 2010; Yim et al., 1985) showing that the doxycycline can reach the choroid plexus (CP). From our transcriptomic analysis, out from 27,164 genes, only 16 where differentially expressed (*Alas2, Bgn, Gypa, Hba-a1, Hbb-b1, Prl, Slc4a1, Snca, Asb1, Mab21l4, Rrp1b, Hba-a2, Btbd2, Xist, Nefh* and *Hbb-bs*). From these genes the significantly DEG where *Hba-a1, Hbb-b1* and *Hba-a2*, all of them related to hemoglobin interaction. An increase in the hemoglobin counts in an organism could be induced by the exacerbated production of red blood cells in the bone marrow. Previously have been shown that haematopoietic progenitor have a higher reprogramming efficiency (Eminli et al., 2009). (Abad et al., 2013) discussed how this reprogramming success and increase in the haemotopoietic progenitors could be the responsible for the teratomas spreading in the different organs through the cyclic induction. As we did not observed major transcriptomic changes in the LVCP we hypothesize that the fact that the choroid plexus epithelial cells (CPECs) has no turnover (McDonald & Green, 1987) might mean that these cells are already in a “senescent” state before the rest of the organs and forbid them to reprogram. A study performed by Browder et al., (2022) to mice carrying a single copy of an OSKM polycistronic cassette and a reverse tetracycline transactivator (rtTA) in a C57BL/6 (B6) genetic background in the gene *Col1a1* evaluated a long-term (from 15-months old to 22 months old and between 12 and 22 months old) and short-term (from 25 to 26 months old) reprogramming treatment by administrating doxycycline (1 ml/mg) in drinking water for 2 days followed by 5 days of withdrawal. They observed that short-term reprogramming of older animals may not work as the cells have already entered a “senescent” state which prevents reprogramming. Also Browder et al., (2022) suggest a hypothesis that some organs as skin and kidney might be more susceptible to reprogramming than other tissue due to different epigenetic status where the CP could be. Another hypothesis of why no transcriptomic changes were observed could be explained by recent studies that evaluate how the expression of Sox2 inhibit the expression of many genes and is recovered once Sox2 is inhibited (Liu et al., 2014). To be able to see transcriptomic changes within groups more time between OSKM induction and sacrifice needs to be evaluated.

Previous work have described the relationship with the LVCP morphology and the CSF production (May et al., 1990; Sakka et al., 2011; Serot et al., 2001, 2003), as we observed morphological changes in the LVCP we decided to look on the 25 most abundantly expressed genes annotated to be secreted factors (Lun et al., 2015) where we did not observed any significant difference on expression between the control and OSKM group.

As no changes in the secreted factors were observed from the transcriptomic analysis, we decided to evaluate the expression of the NKCC1 protein, which is a Na-K-Cl cotransporter responsible for 50% of CSF production (Steffensen et al., 2018). A significant increase in NKCC1 protein expression was observed in the CPECs even though we did not observe this increase in the relative mRNA NKCC1 expression from the bulk RNA-seq analysis. The translation rate is mainly regulated by the mRNA sequence but it could also be regulated by the presence of micro-RNA or long noncoding RNA (lncRNA) (Khatir et al., 2023) which are not evaluated in the transcriptomic analysis here performed. Specific micro RNA (miRNA) have been identified and used to increase reprogramming efficiency as they are able to target multiple pathways without needing a specific amount transcription factors to achieve it (Omole & Fakoya, 2018). According to (Uniprot) there is one identified isoform from the coding gene of NKCC1 (*Slc12a2*) which was not considered in the bioinformatics analysis either. To further understand the dysregulation between the transcription and translation of NKCC1 deeper transcriptomic and proteomic analysis need to be performed.

With a higher production of CSF due to an increase in the protein expression of NKCC1, we evaluated the reaction of the ventricular-subventricular zone (V-SVZ) where is the largest germinal zone in the adult mouse brain (Mizrak et al., 2019; Silva-Vargas et al., 2016). The lateral walls for the ventricles hosts the neural stem cells (NSCs) in the adult brain. Two types of NSC have been described as quiescent (qNSC) and actively dividing (aNSC) states (Chaker et al., 2016). Sox2 is a transcription factor that is expressed in the proliferating NSCs and during the developing and mature brain (Liu et al., 2014) and it is also part of the Yamanaka factors induced to achieve reprogramming. The increase of Sox2 in the V-SVZ ependymal cells indicate thar rejuvenation is at stake in the brain too. To evaluate is this expression is caused by the CSF secreted by the CP or through different pathways within the brain a broader study targeting the signaling cascade the reprogramming causes in the brain will need to be performed.

## Figure legends

**Supplemental Figure 1.**
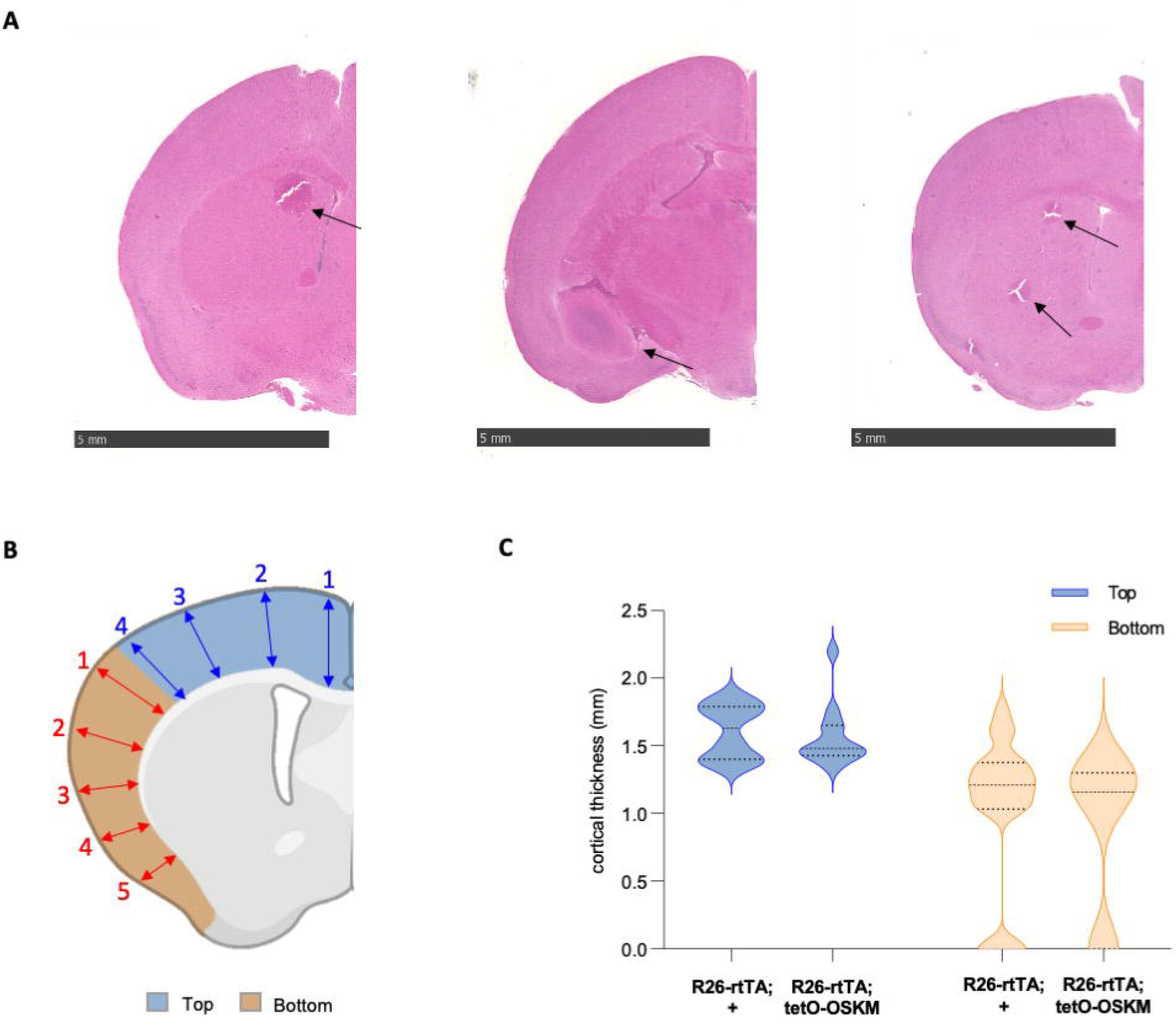

