## Supplemental information for "Impact of in vivo cyclic reprogramming on the choroid plexus"

### Supplemental Figure legends

**Supplemental Figure 1. Effects of global cyclic OSKM induction on the brain.**

**(A)** Representative coronal cuts of whole brain of R26-rtTA;tetO-OSKM mice stained with hematoxylin and eosin. Teratoma is indicated with a black arrow. Scale bar: 5 mm. **(B)** Schematic depicting a half coronal brain cut with the two areas where cortical thickness was measured. The blue area represents the "top" of the cerebral cortex. The orange area represents the "bottom" of the cerebral cortex. The thickness of the top and bottom areas was measured at different locations as indicated by numbered arrows. **(C)** Cortical thickness of the top and bottom areas was measured on coronal cuts of whole brain of R26-rtTA;+ control mice and R26-rtTA;tetO-OSKM mice.

### Supplemental Figure 1

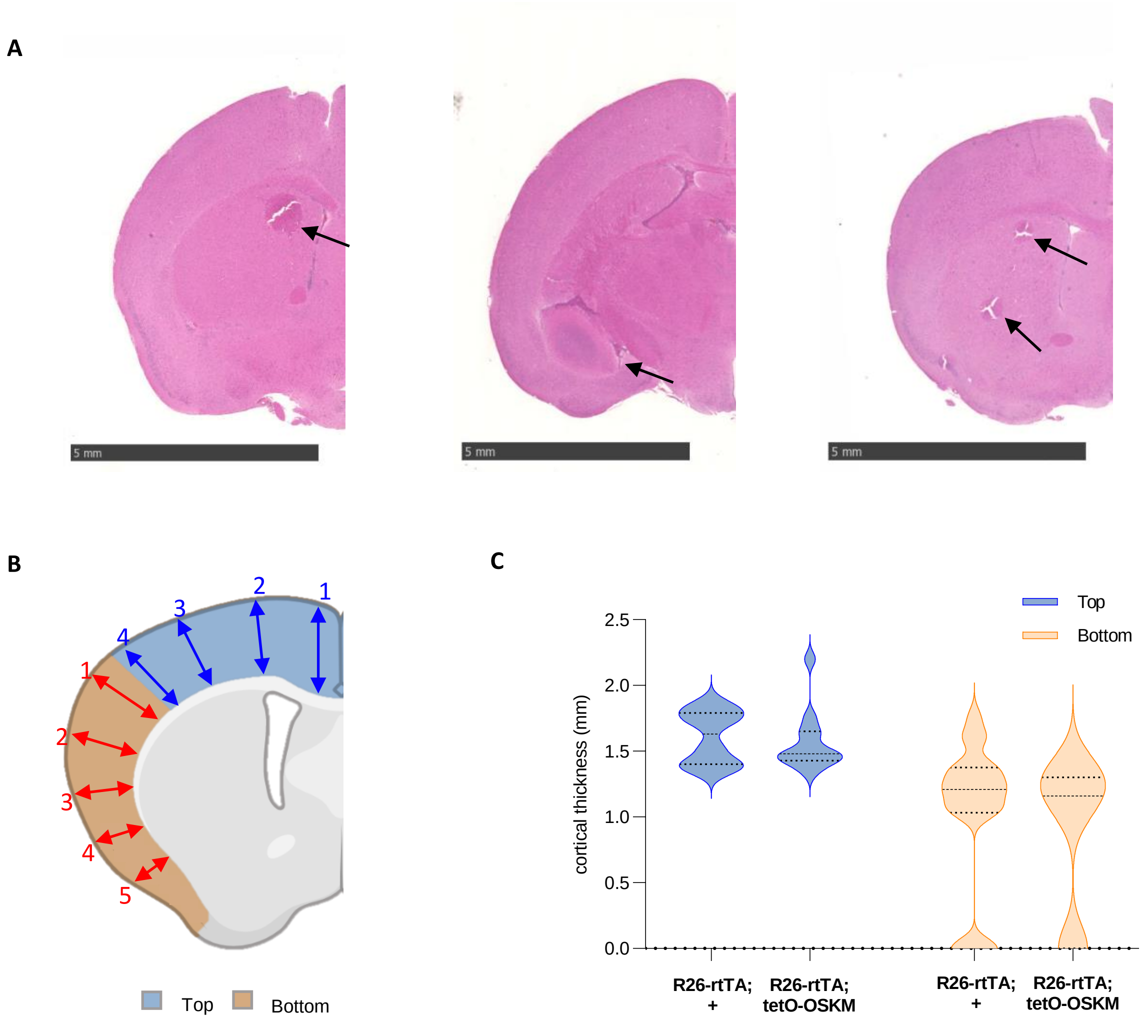
